# Transcription network of SLC7A11 (xCT) in colon cancer provides clinical targets for metabolic regulation and cell proliferation

**DOI:** 10.1101/2024.06.03.597098

**Authors:** Keren Zohar, Thomas Wartmann, Marco Strecker, Maximilian Doelling, Mihailo Andric, Wenjie Shi, Roland S Croner, Or Kakhlon, Yue Zhao, Ulf D Kahlert, Michal Linial

## Abstract

Colorectal cancer (CRC) represents the third leading cause of cancer-related deaths. Knowledge covering diverse cellular and molecular data from individual patients has become valuable for diagnosis, prognosis, and treatment selection. Here, we present in-depth comparative RNA-seq analysis of 32 CRC patients pairing tumor and healthy tissues (total of 73 samples). Strict thresholds for differential expression genes (DEG) analysis revealed an interconnection between nutrients, metabolic program, and cell cycle pathways. Among the upregulated DEGs, we focused on the Xc-system, composed of the proteins from SLC7A11 (xCT) and SLC3A2 genes, along with several interacting genes. To assess the oncogenic potency of the Xc- system in a cellular setting, we applied a knowledge-based approach, analyzing gene perturbations from CRISPR screens. The study focused on a set of 27 co-dependent genes that were strongly correlated with the fitness of SLC7A11 and SLC3A2 across many cell types. Alterations in these genes in 13 large-scale studies (e.g., by mutations and copy number variation) were found to enhance overall survival and progression-free survival in CRC patients. In agreement, the overexpression of these genes in cancer cells drives cancer progression by allowing effective management of the redox level, induction of stress response mechanisms, and most notably, enhanced activity of ion/amino acid transporters, and enzymes acting in de novo nucleotide synthesis. We also highlight the positive correlation between the Xc- system gene expression level, patient responsiveness to different chemotherapy treatments, and immune cell infiltration (*e.g.,* myeloid-derived suppressor cells) in CRC tumors as a measure for their immunosuppressive activity. This study illustrates that knowledge-based interpretation by synthesizing multiple layers of data leads to functional and mechanistic insights into the role of SLC7A11 and its associated genes in CRC tumorigenesis and therapeutics.

## Introduction

Colorectal cancer (CRC) represents 10% of cancer cases globally. It is the third most prevalent cancer and the third leading cause of cancer-related deaths in the USA. Having access to extensive data sets from cancer patients enables the evaluation of crucial predictive biomarkers needed for optimizing treatment choices. Moreover, earlier diagnosis is crucial for improved survival. At present, diagnosis and choice of optimal treatments are based on integration of clinical features (e.g., age, family history, tumor location, size and TNM staging) (Vega et al. 2015). For higher precision, genetic alterations are tested, in particular evidence for recurrent mutations (Testa et al. 2018). While noninvasive tests (e.g., blood and stool tests) show promise for improved detection, the lack of mechanistic or cellular interpretation limits their use (Zygulska and Pierzchalski 2022). The primary treatment for most CRC patients is surgery, with chemotherapy being administered in cases of advanced disease (Biller and Schrag 2021). The multi-omics approach that includes diverse cellular and molecular data from individual patients at a large-scale has become a valuable component in cancer research, influencing CRC diagnosis, prognosis, and treatment selection (Menyhárt and Győrffy 2021).

Ample studies have shown the importance of pathological measures that distinguish between normal and tumor tissues based on histological and rich clinical criteria (Roseweir et al. 2017). In recent years, rich data from hundreds of patients were compiled in cancer portals (e.g., TCGA, GDC, cBioPortal), providing rich omics data such as gene expression and epigenetic profiling which had been successfully used in prognosis of various malignant tumors (Chen et al. 2022). In the case of CRC, there is an urgent need to identify precise prognostic factors to identify patients who would benefit most from proposed treatments. In terms of preferred treatments, agents that lead to DNA synthesis disruption and eventually cell death are commonly used (e.g., 5-fluorouridine or fluoropyrimidines) (Mármol et al. 2017). While other drugs are available (e.g., oxaliplatin, fluoropyrimidine, irinotecan), there is little evidence that can match the beneficial use of any of the available agent to specific patients based on their detailed molecular profiles (Koncina et al. 2020). The analysis of healthy and tumor samples from the same patient is expected to improve prognostic accuracy in CRC patients (Barrier et al. 2005).

The prognosis of CRC patients with advanced tumors remains poorly understood. Tumor-infiltrating immune cells were proposed to impact cancer progression, treatment response, and ultimately patients’ survival and therapy efficacy (Zheng et al. 2022). For example, the presence of tumor-associated neutrophils (TANs), tumor-associated macrophages (TAMs), and myeloid-derived suppressor cells (MDSCs) has been linked to worsened prognosis in CRC and other cancer types (Condamine et al. 2015; Parcesepe et al. 2016). Comprehensive analytical algorithms use information from large-scale studies *(e.g.,* TCGA) to assess sample purity and the correlation of immune cell infiltration with gene expression as a means to provide insights into the predictive value for treatment decisions for CRC patients. Moreover, studying tumor cells’ metabolic demands, and their capacity to cope with stress, along with characterizing the cancer immune microenvironment, can benefit therapy precision (Li et al. 2020; Li et al. 2021).

In any living system, the accessibility of amino acids is crucial for energy production, translation efficiency, and redox homeostasis (Vučetić et al. 2017). The dysregulation of amino acid transporters in tumor cells complies with the increased metabolic and translational needs (Zhu and Thompson 2019). Cancer cells require large amounts of cysteine and glutathione (GSH) to neutralize the increased intracellular reactive oxygen species (ROS). Cysteine plays a major role in maintaining antioxidant defense in cancer cells, highlighting its significance in cellular redox balance. The main challenge is that while the intracellular environment favors a reducing state, the extracellular environment is strongly oxidizing, leading to the rapid oxidation of cysteine to cystine (Daher et al. 2020).

Cancer cells, facing high oxidative stress, struggle to meet their demand for cysteine (Bonifácio et al. 2021). Cystine starvation induces cell death that can be rescued by antioxidants. Most cancer cells rely on the Xc- heterodimeric amino acid transporters system, consists of SLC7A11 (xCT) and SLC3A2 (heavy chain 4F2hc), which imports cystine for glutathione synthesis (Koppula et al. 2021). The Xc- system consists of chloride-dependent anionic L-cystine/L-glutamate antiporter on the cell surface, which mediates the uptake of extracellular cystine in exchange for intracellular glutamate (Lin et al. 2020). Briefly, Xc- system imports cystines (the oxidized form of cysteine) that ultimately serve as precursors for reduced glutathione (GSH) synthesis. Tripeptide GSH synthesis involves two enzymatic steps, starting with the condensation of cysteine and glutamate into γ-glutamyl-L-cysteine by glutamate cysteine ligase (GCL) followed by adding glycine to form GSH by GSH synthase (GS) (Lin et al. 2020). GSH is involved in several vital cellular functions, including detoxification, maintaining intracellular redox balance, reducing hydrogen peroxide and other oxygen radicals, and serving as thiol donor to proteins. While there are other transporters that can partially compensate for a failure in SLC7A11, it remains the major route for transporting cystine in cancer cells (Parker et al. 2021). Similarly, in cancer stem cells (CSCs), CD44 variant isoform (CD44v) can interact and stabilize SLC7A11 on the cell surface (Jyotsana et al. 2022).

While SLC7A11 was identified 40 years ago, details on its expression regulation in cancer cells in view of metabolic load, redox status, and tumor microenvironment (TME) remain incomplete. The nutrient dependency of cancer cells generally requires the increased function of SLC7A11. A high expression of SLC7A11 leads to a reduction of oxidative stress in some oncogenic KRAS-mutant cancers and thus maintains cancer progression (Koppula et al. 2021). In most cases, the elevated SLC7A11 expression is related to a low survival rate. This was validated in the cases of pancreatic ductal adenocarcinoma (PDAC), colorectal adenocarcinoma (COAD) and lung adenocarcinoma (LUAD). In contrast, SLC7A11 knockdown leads to an oxidized redox status and an increase in intracellular ROS levels. Ultimately, such stress may inhibit tumor invasion.

The discovery of ferroptosis, a form of regulated cell death induced by iron-dependent lipid peroxide accumulation, through blocking cystine uptake, further highlights the importance of the cystine transport system for cell survival. Pharmacologic blockade of SLC7A11 induces ferroptotic cell death (Dixon et al. 2012). Studies show that SLC7A11 mediated cystine uptake is essential in suppressing ferroptosis and promoting cell survival under oxidative stress. Thus, the regulation of the Xc- system is an attractive target for cancer therapy (Lei et al. 2022). Nevertheless, there are discrepancies between the pro- and anti-tumorigenic activities of SLC7A11 when utilizing a simplified setting of cell culture versus in vivo models (Li et al. 2022). The amounts and activity of SLC7A11 in the cell membrane are strongly regulated at the transcription level, but also respond to translational and post-translational regulation (Lee and Roh 2022). SLC7A11 expression is induced by ATF4 under amino acid deprivation, which is essential for cells to survive under conditions of cystine starvation-induced ferroptosis (Zou et al. 2024). Understanding how these mechanisms modulate SLC7A11 can provide insight into therapeutic targets for cancer treatment.

In this study, we focus on the transcriptional profiles (mRNAs) from 32 colon cancer patients, each analyzed by comparing its tumor to the healthy tissue. We identified strong upregulated gene sets that signify mitotic cell signature from colon, cell cycle G2M checkpoints and an additional network of dysregulated transporters leading to a metabolic burden. We focused on SLC7A11 and its functional network as an integrator of colon cancer progression. Using functional CRISPR cellular fitness analysis and survival data from large resources of colon cancer, we identified genes carrying clinically relevant properties. We present an exhaustive bioinformatic analysis to explore the impact of the Xc-system on therapy responsiveness and the tumor composition of immune cells. We illustrate the importance of a multilayer analysis, initiated from detailed transcriptional tissue profiling, in exposing overlooked cellular processes and targets for improving clinical and therapeutic management of CRC.

## Methods

### RNA-seq analyses of 32 CRC patients

The mRNA expression levels of all genes (coding and non-coding) were determined by pairwise analysis of cancerous and healthy tissues obtained from the same patient. Total of 32 patients were analyzed with 73 deep sequencing results. Altogether, there were 36 samples marked as tumor (T) and 37 samples marked as healthy (H). Each participant provided at least one sample for T and H. For four participants the number of samples was higher.

Ethical approval to conduct this study was granted by the ethics committee of the medical faculty of Magdeburg (33/01, amendment 43/14). Next-generation sequencing was conducted in contract-based cooperation at the genome analytics lab at Helmholtz-Center for Infection Research (HZI) Brunswick, Germany.

### Analyses of CRC patients from public resources

The Limma R package (Ver 4.2.0) was used for differential expression analysis with adjusted p-value of 1e-20 (for pair-wise analysis) as significance threshold. We have applied GEPIA2 database that covers the data from The Cancer Genome Atlas (TCGA) and Genotype-Tissue Expression (GTEx) (Consortium 2013). Box plot, violin plot, and scatter plot for selected DEG were drawn by the TCGA and GTEx visualization website GEPIA2 (Tang et al. 2019).

### Colon cell type

Analysis of bulk RNA-seq datasets from 15 human organs including colon produced a cell type enrichment prediction atlas for all coding genes. The initial data is extracted from GTEx. The identity profiles across tissue types revealed 12 types of cells in colon by the Human Proteome Atlas (HPA; (Thul and Lindskog 2018)). We have applied the resource for understanding the DEG from colon cancer. The 12 colon cells cover 1918 genes (559, 622 and 737 that are labelled as very high, high and moderate enriched genes, respectively). We performed the analysis for 7 main colon cell types Colon enterocytes (369 genes), Colon enteroendocrine cells (338 genes), Enteric glia cells (240 genes), Mitotic cells in Colon) 85 genes), Endothelial cells (219 genes), Smooth muscle cells (166 genes), Fibroblasts (42 genes). Additionally, there are 5 types of immunological cells of the colon that are specified by their enriched genes: Macrophages (143), Neutrophils (65), Mast cells (29), T-cells (108) and Plasma cells (114).

### Bioinformatics tools and statistics

#### Statistically significance

Paired statistics for 2-group analysis was based on 2-taileed t-test. Statistical significance was also computed using Mann-Whitney. Kruskal-Wallis tests used in single-variable comparisons with more than 2 groups. Differences with *p* <0.05 were regarded as statistically significant (unless mentioned otherwise). False Discovery Rate (FDR) was computed using the Benjamini-Hochberg method. Hypergeometric test was used to assess the p-value of overlapping gene sets.

#### Signature gene set

The database of gene sets from the Molecular Signatures Database (MsigDB) allows to test gene set enrichment and it includes about 10,000 sets covering diverse biological processes and diseases. A collection of hallmark gene sets is a set of 50 main processes in cells with expert curation with about 200 genes included in each hallmark set (Liberzon et al. 2015).

#### Gene expression density plot

Conducted using RNA-seq data from TCGA, combined with the Therapeutically Applicable Research to Generate Effective Treatments (TARGET), and the GTEx repositories using TNMplot (Bartha and Gyorffy 2021).

#### Enrichment tests for cancer hallmarks

Testing overrepresentation analysis by slice representation. The different colored slices indicating the hallmarks (total of 10) that are significant (using the adjusted p < 0.05 as a threshold). The analysis used 6763 genes that are associated with any of the hallmarks as a reference set.

### Expression of immune cells

We used DICE resource that displays the gene expression trend for 13 immune cells in their naïve and activated states. DICE identified cis-eQTLs for 61% of all protein-coding genes expressed in these cell types (Schmiedel et al. 2018). For purity and infiltration of immune cells to the tumor sample, we used the TIME2.0 (Tumor immune estimation) resource that applied correlation tests or any gene against 22 immune cell types. The TIDE platform report on any gene (or gene sets) across over 33K samples in over 180 tumor cohorts (including TCGA) for the T cell dysfunction and exclusion signatures associated with it (Fu et al. 2020).

### CRISPR-Cas9 cell line screening

We used the pre-calculated correlation of dependency from DepMap using CRISPR/Cas9 (Dempster et al. 2019). CRISPR-Cas9 and RNAi-based knockout are reported for 19,144 genes across 1206 cell lines (primary and established) and providing knockout fitness scores (measured 14 days after transfection) and control metrices for the calculation of probabilities of dependency across cell lines that are divided by their origin and lineage. Dependencies enriched in COAD were precalculated for ∼1800 genes (identified by DepMap CRISPR-Cas9 project using the Public 23Q4+Score, Chronos resource). Expanded collection of cell lines and cancer types is available in iCSDB (Choi et al. 2021) that combined DepMap (Public 20Q2) and BioGRID ORCS (Ver. 1.0.4) large scale CRISPR data. We search for genes with correlated knockout fitness (called ‘co-dependent’). A loss of fitness and a negative log fold change in the average representation of the relevant targeted sequence relative to plasmid are indicative for the gene being essential.

### Predictive analysis by gene expression level

KM Plot and ROC Plotter (Fekete and Győrffy 2019) were used to identify gene expression-based predictive biomarkers for CRC that compiled publicly available datasets. By integrating gene expression data (RNA-seq and Chip-Seq) with chemotherapy, almost 20,000 genes can be tested. A link of gene expression and therapy response using transcriptome-level CRC data generates a ROC plot with detailed statistics on relevance of any gene to therapy and clinical response (Fekete and Gyorffy 2023). In addition, we activated a platform for validating predictive biomarkers in cell lines (>1200 cell lines, based on 4 resources including DepMap). The expression of genes in cells with and without drug treatment are presented by the average response per each cell collection.

## Results

### Pairwise analysis of samples from colon cancer patients

We have analyzed 32 colon cancer patients with 73 datasets. Each patient contributed at least two samples, one from the tumor and another one from the unaffected neighboring tissue. A few patients, had more samples (3-6 each). All samples were subjected to deep sequencing for mRNA profiling (see Methods).

**Fig. 1A** shows the unsupervised partition of all 73 samples labelled tumor and healthy (T and H, respectively). The dendrogram shows a clear partition of all samples into two main branches. The tumor (T) branch is 100% consistent (purple, 31 samples), and the second major branch is mostly composed of healthy samples (88% of 42 samples, orange), only 5 T-labelled samples clustered with the H-samples. **Fig. 1B** shows that dimensional reduction by principal component analysis (PCA) supports a successful partition of T and H samples with 37% of the total variance explained by PC1 and PC2. The PCA used the top 1000 ranked differentially expressed genes (DEG). Similar successful partition of T and H was achieved by PCA whose input consisted of the entire (*i.e.,* not only DEG) RNA-seq profiles (total 13,682 identified transcripts).

**Figure 1.**
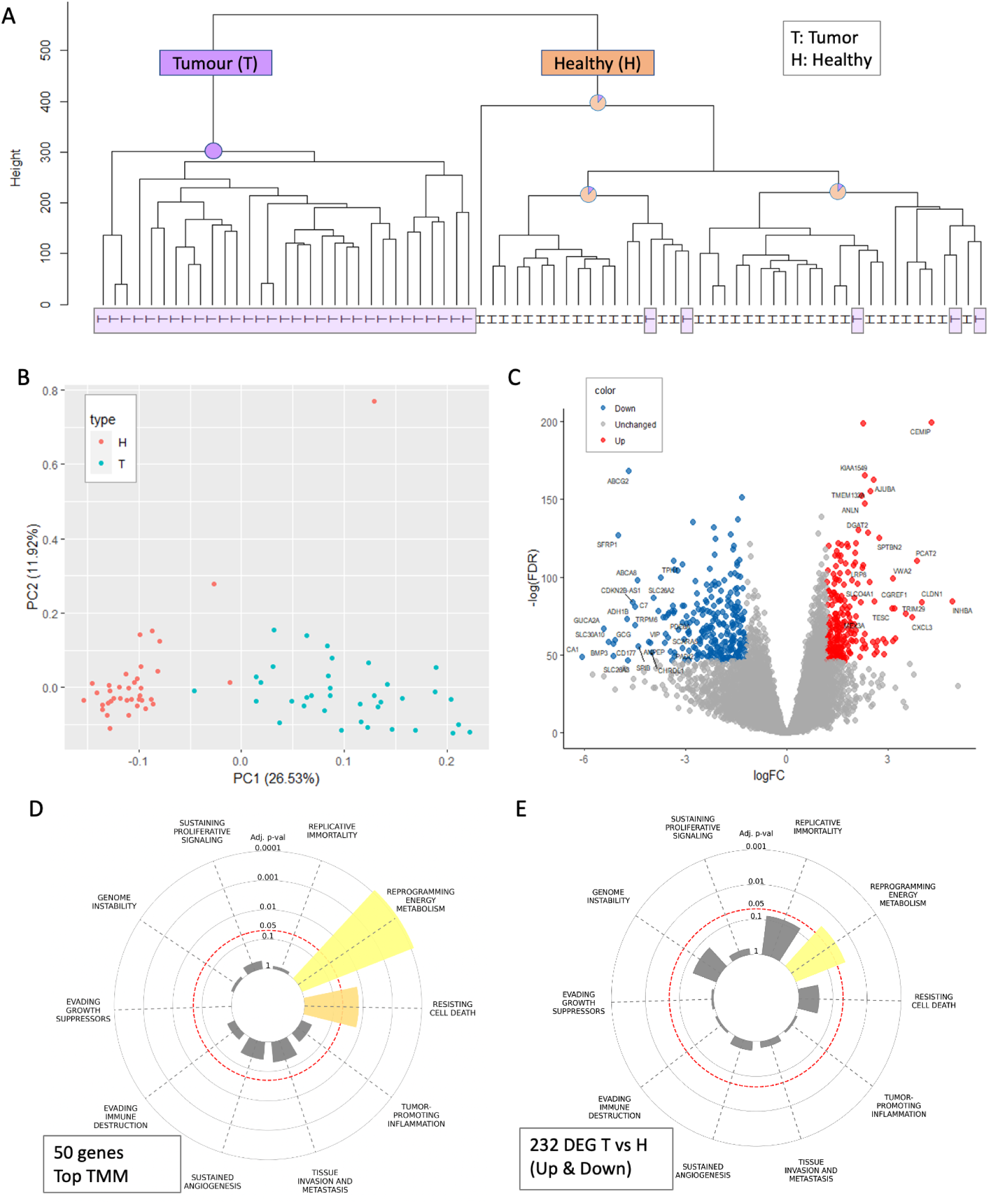
Analysis of the mRNA profiles from 32 colon cancer patients. **(A)** Unsupervised dendrogram of 73 samples from 32 participants. The main nodes are indicated by their purity for tumor (T) and healthy (H) colored purple and orange, respectively. The T samples of the dendrogram tree are highlighted with light purple background. **(B)** PCA for 73 samples, based on the top 1000 differentially expressed genes (DEG) colored by T and H with red and blue, respectively. The variance explained are indicated for PC1 and PC2. **(C)** Volcano plot representation of DEG analysis from RNA-seq of T versus H for the samples described in A. Red and blue points mark the genes with significantly increased or decreased expression in T relative to H, respectively. Representative significant genes are indicated. **(D)** Top 50 expressing genes (RNA-seq, normalized by trimmed mean of the M-values (TMM) tested for enrichment for any of the 10 cancer hallmarks. **(E)** DEG (up and down; 323 genes). The significantly enriched hallmarks are colored.

Next, we analyzed the DEG from all colon cancer patients. Each patient was analyzed with respect to their own healthy-labelled sample. Single samples from the tumor and healthy tissue were normalized and compared internally (according to the number of samples available). Altogether, we performed global analyses of 32 pre-analysis patients to confirm high statistical significance and a minimal fold change threshold per gene. Specifically, the analysis was restricted to genes with a minimal statistics of FDR p-value <1e-20, with a minimal average expression of 10 counts per million (CPM) and limited to coding genes (*i.e.,* 92% of all mapped transcripts). Such filtration reduced the 13,682 unique gene transcripts to 9,045 genes that were further analyzed (**Fig. 1C**).

We tested the results of the RNA-seq analysis to identify a signature for any of the 10 known cancer hallmarks (Hanahan 2022). The highly expressed genes already identified significant hallmarks such as ‘reprogramming energy metabolism’ (p-value <1e-06) and ‘resisting cell death’ (adjusted p-value 1.4e-02; **Fig. 1D**).

For clinical relevance, it is essential to focus on consistent expression difference in T to H samples. To this end, we reanalyzed DEG at a relaxed threshold. For enrichment of cancer hallmarks DEG were selected with FDR <1e-20, a minimal fold change of 2.3 (*i.e.,* log(FC) >|1.2|), and a minimal average expression of 10 CPM (Supplementary **Table S1**). The signature for ‘reprogramming energy metabolism’ remained significant (adjusted p-value =2.7e-02) (**Fig. 1E**). We concluded that among the identified DEG from the CRC patients, a signature of metabolic programming dominated.

### Colon cancer DEG reveals hallmarks of cell cycle and metabolic program

We further tested the enrichment of DEGs (323 genes: upregulated: 130; downregulated: 193) with respect to the predetermined 50 cell hallmark sets (about 200 genes each, see Methods).

Manually selected 50 gene sets cover major cellular biological processes (see Methods). **Table 1** lists the most enriched sets (Adjusted p-value <1e-06). Several observations can be made based on the results in **Table 1**. Firstly, the stronger enrichment is for a set of genes encoding cell-cycle related targets of E2F transcription factor which is exclusively composed of upregulated genes. Moreover, for most significantly enriched cell hallmarks, the gene set labelled consists of genes with the same trend, either up- or downregulated genes. The only exception is the hallmark called ‘estrogen response-late’ that shows a mixture of up- and down-regulated genes (notably, it was primarily based on breast cancer data).

**Table 1.**
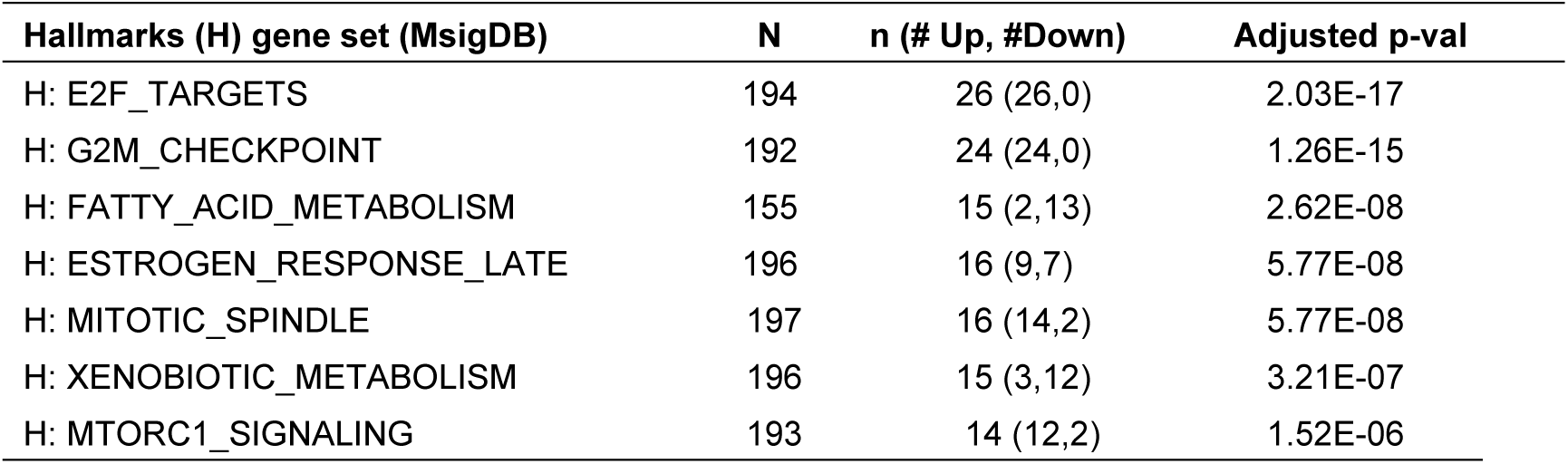
Enrichment analysis of 323 DEG for the 50 gene set of MSigDB hallmarks.

The results from **Table 1** can be broadly classified into two larger themes: cell cycle-related (*e.g.,* mitotic spindle, G2M checkpoint and genes encoding cell cycle E2F) and nutrients and metabolic programs (*e.g.,* fatty acids synthesizing, mTOR signaling, and genes involved in processing of drugs and other xenobiotics). **Fig. 2A** tests the overlap of the hallmark sets that belong to these main themes. With 20 genes overlapping the cell cycle, 8 genes signifying mTOR signaling and only 4 genes overlap all gene sets (marked in **Fig. 2B**, by stars). The connectivity of the upregulated genes identifies a dominant network of cell cycle G2M checkpoint (**Fig. 2B**, red) and a smaller cluster of cell membrane transporters including SLC11A7 that match mostly genes that specify nutrient and metabolic management in cells (**Fig. 2B**, green).

**Figure 2.**
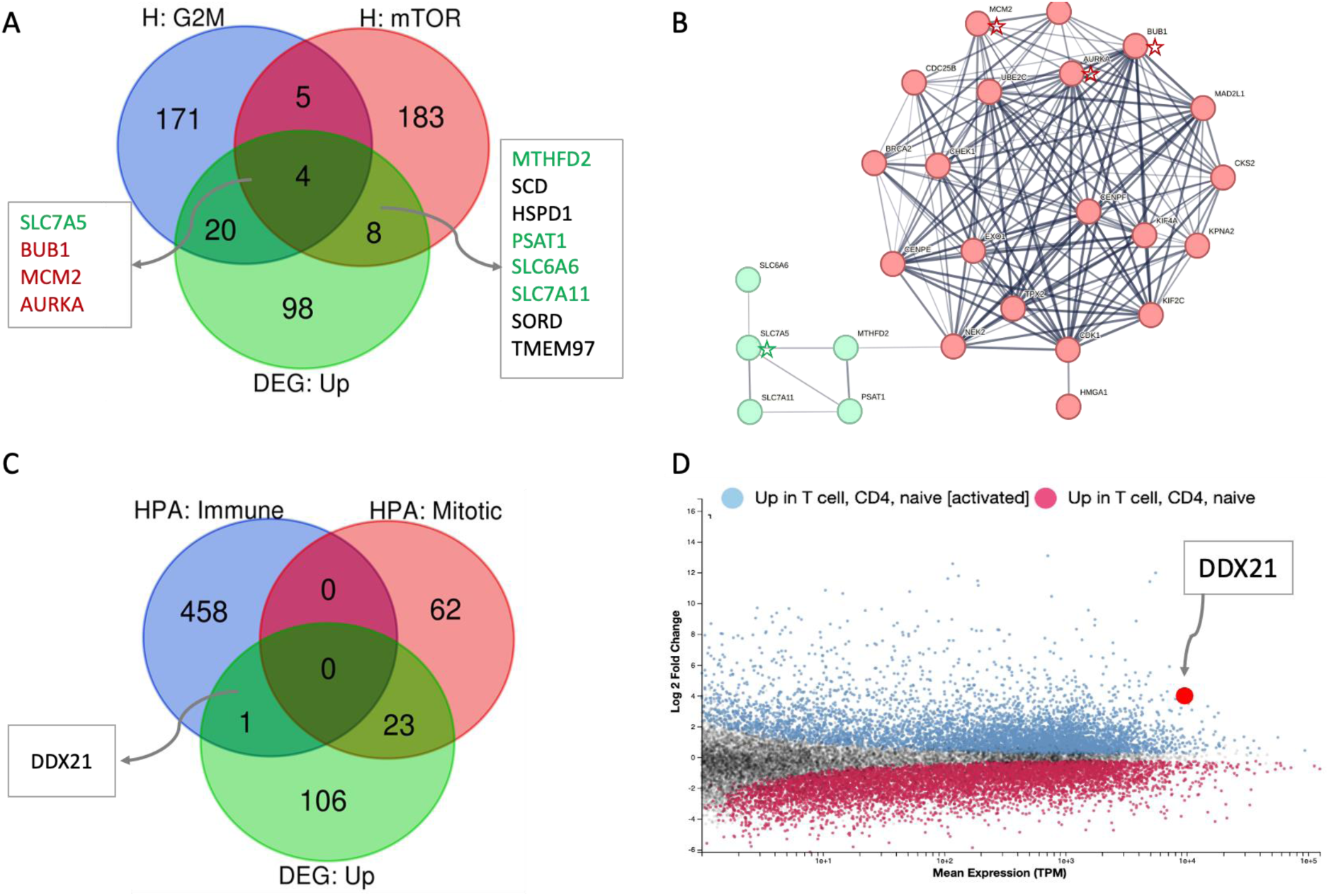
Overlap of upregulated DEG with the dominant hallmarks cellular processes. **(A)** Venn diagram of the upregulated DEG (130 genes) and the hallmark sets of ‘cell cycle G2M checkpoint’ and ‘mTORC1 signaling’. The overlap genes are listed. **(B)** STRING network of the overlap DEG from (A) with the G2M and mTOR sets (total 20, 8 and 4 genes). STRING confidence score >0.4 shows the PPI connected gene network. The genes shared by all three sets (4 genes, see A) are indicated by stars. The cluster in green includes overlapping genes. Gene names are shown by a green font (in A). **(C)** Venn diagram of the colon cell types lists for the immune related unified set and the colon mitotic cell types. **(D**) DEG analysis of naïve CD4 T-cells and following their activation (see Methods). The overexpression of DDX21 is shown by the arrow. DEG with FDR at a threshold of <1e-20 are colored gray. X-axis shows the calculated mean gene expression (by TPM) and x-axis the log2(FC). Following activation, gene expression of DDX21 is ∼16 fold higher than the naïve CD4 T cell basal level.

### The upregulated genes strongly identified subpopulation of mitotic cell types

The colon is a complex tissue composed of numerous cell types. The composition of the cell types in colon was determined from single cell and bulk data analyses (see Methods). We tested the set of upregulated genes (total 130, Supplementary **Table S1**) with respect to the 12 characterized cell types that are signified by enriched gene sets. Among these 12 cell types, the 5 immunological cells (see Methods) were excluded no overlapping genes were identified. We tested the upregulated DEG for each of the other 7 main colon cell types. A significant overlap was found only to mitotic cells, with 23 DEG overlapping 85 mitotic cell enriched genes (enrichment p-value 3.5e-05). The PPI network of the 23 genes is highly connected with an average node degree of 14.1 and a PPI enrichment p-value <1.0e-16. Among these genes are kinesin-like proteins that act in chromatid segregation (KIF2C, KIF14, KIF20A), numerous genes that participate in cell cycle via DNA repair mechanism (RAD51AP1, EXO1, BRCA2) and genes involved in the checkpoint controls for ensuring DNA replication (TPX2, BUB1, NUF2, CDC6). A full list of all genes by their cell types are listed in Supplementary **Table S2**.

We then asked whether there is evidence for colon enriched immune cell signature among upregulated DEG. We compiled a colon-centric immune enriched set by unifying all 5 immune cell types (total 459 genes, see Methods). Interestingly, DDX21 (FDR 5.30e-46) is the only gene that matched the unified colon immune-related gene set (**Fig. 2C)**. In the context of colon cancer, knockdown of DDX21 inhibited cell growth by activating CDK1, which was also identified among upregulated genes overlapping the mitotic signature. DDX21 was postulated to mediate this effect via chromatin modulation of the CDK1 promoter (Lu et al. 2022). The nucleolar activity of DDX21 is in rRNA processing and ribosome biogenesis. Interestingly, following activation of T cells, the gene is upregulated to an extreme level (**Fig. 2D)**.

### Enrichment of extracellular and plasma membrane regions among the strongest DEG

We then tested the possibility of identifying tumor versus healthy genes and focused on the subset of genes showing the most extreme differential expression signals. To this end, we analyzed a subset of 84 DEGs with fold change (FC) of >|5|. While this is an arbitrary threshold, it captures the maximally responding DEG group.

**Fig. 3A** shows that at this threshold, 80% of the genes were downregulated and only 20% were upregulated. **Fig. 3B** shows a connectivity map of these DEG (with at least 2 gene connections; STRING confidence score of 0.5). Most connected genes (70%) were assigned with either extracellular regions (GO annotation of cellular component, p-value =3e-06; colored red) or plasma membrane region (p-value =0.003; colored blue). These significant findings suggest that in colon cancer, the most differentially expressed genes probably act through an extracellular communication, and potentially act in transport and signaling at the plasma membrane.

**Figure 3.**
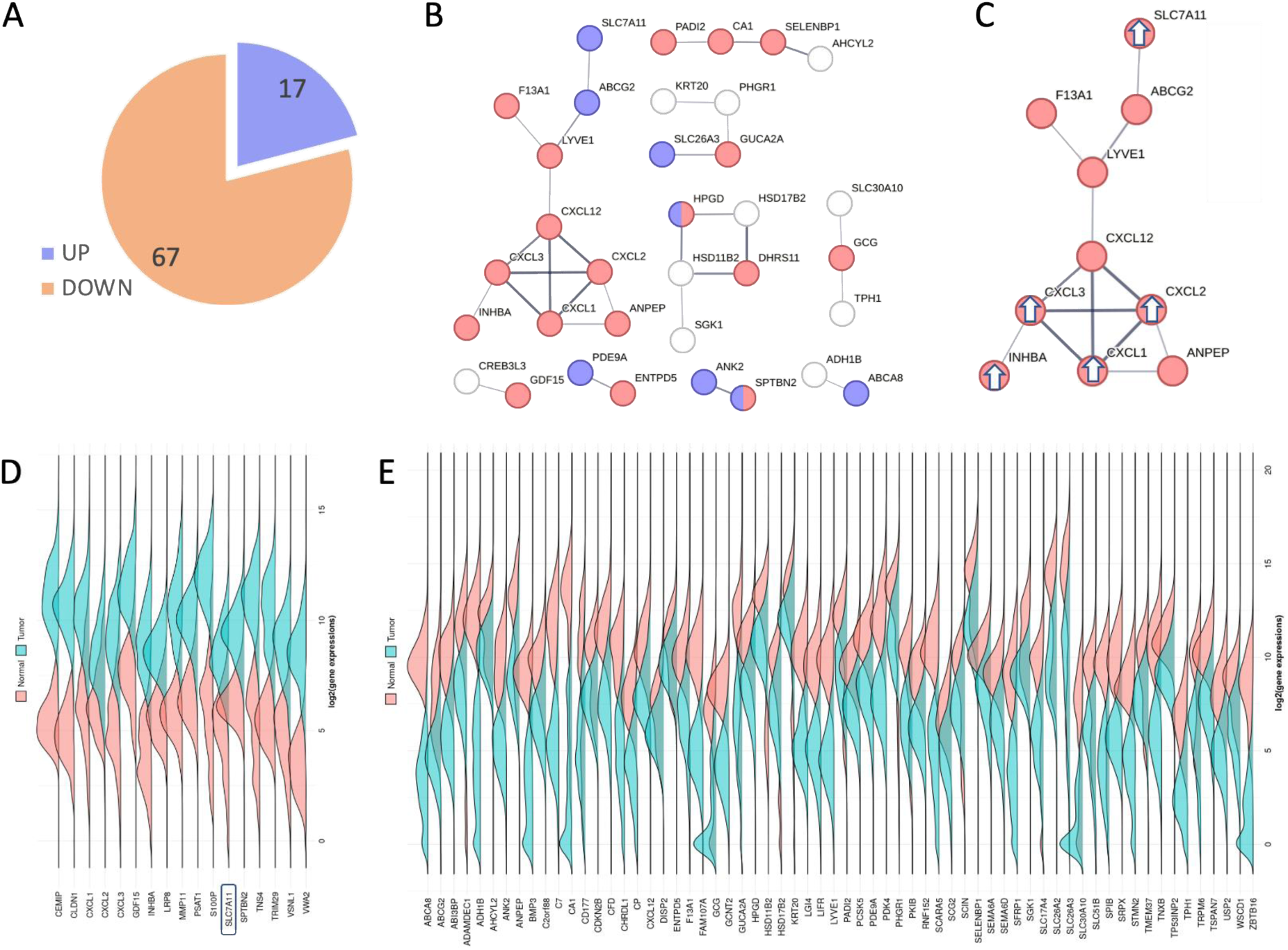
Biological insight from DEGs with extreme fold change (FC>|5|) of colon cancer samples. **(A)** Pie chart of 84 DEG with FC >|5|, partitioned to up and downregulated genes (20% and 80%, respectively). (**B)** STRING based network (confidence threshold 0.5). Only confident connected genes are shown. Colored are genes that are annotated by GO annotation of cellular component as extracellular and plasma membrane regions (red and blue, respectively). (**C**) The largest connected component from B. DEG that were upregulated are marked with white arrows. The other nodes are genes that were downregulated. **(D)** Density plot analysis of the 17 upregulated DEG (alphabetic order). Expression density plots of healthy and tumor samples are in pink and blue, respectively. The SLC7A11 gene is marked. **(E)** Density plot analysis of the 67 downregulated DEG (alphabetic order) for COAD.

**Fig. 3C** shows the largest connected component (10 genes, STRING interaction as in **Fig 3B**). This 10-node subnetwork is the only one that included upregulated genes (marked with arrows), the other subgraphs include downregulated genes. The genes include SLC7A11 transporter, genes that function in mitochondria and a set of secreted cytokines. Notably, among the 84 DEG, cell membrane transporters were overrepresented with 6 genes that belong to the solute carrier family (SLC genes), and 2 belong to the mitochondrial ABC transporters. Only SLC7A11 was strongly upregulated while the rest of the transporter encoding genes were strongly downregulated.

To validate that the DEG from our study are in agreement with the large-scale available data from available cancer resources, we performed density plot analysis for the upregulated **(Fig. 3D**) and downregulated (**Fig. 3E**) genes. The results show that there is a complete agreement in the list of all 84 DEGs (Supplementary **Table S1**) regarding the expression level trends in healthy and tumor samples of our cohort.

### SLC7A11 and its interactors exhibit a coordinated upregulated expression in CRC

**Fig. 4A** lists the validated set of SLC7A11 interactors across multiple tissues (total 16 genes; STRING confidence score 0.7). SLC7A11 interactors display its strong connection to SLC3A2 and CD44 (consistent with the Xc- system) and to other 6 SLC family transporters. In addition, SLC7A11 acts as a hub to cellular metabolic genes. Examples include OTUB1, a specific deubiquitylating enzyme with a cysteine protease activity, BECN1 (Beclin 1) that regulates vesicle-trafficking processes, autophagy, and apoptosis. A number of the SLC7A11 knowledge-based network act under starvation, oxidation and ER stress (*i.e.,* ATF4, NFE2L2, GTX4). The transcription factor ATF4 acts to induce various amino acid transporters and enzymes that determine the metabolic state of cells (including redox balance, autophagy, energy production, and nucleotide synthesis). Other core genes are directly associated with cancer progression (TP53 and EGFR) that drive cell migration, differentiation and cell growth (**Fig. 4A**).

**Figure 4.**
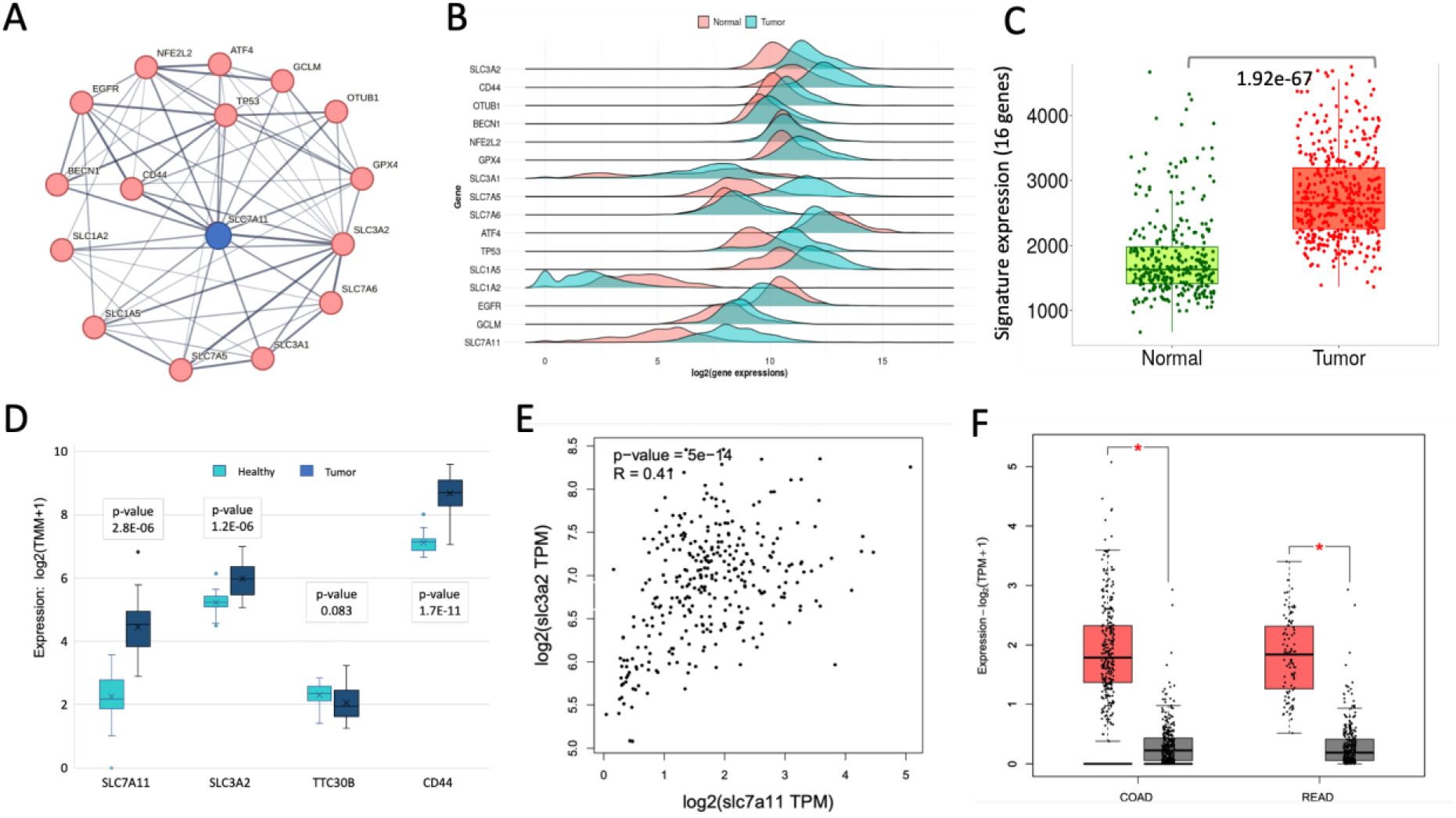
Signature of SLC7A11 network in colon cancer. **(A)** Interacting core genes centered by SLC7A11 according to STRING (confidence score >0.7), limited to the most significant 15 additional genes. **(B)** Density plots for 16 genes from (A). Healthy and tumor samples are marked in pink and blue, respectively. (**C)** Box plot for the signature of all 16 listed genes for healthy (normal, green) and tumor (red). Each dot represents a unified datapoint for a COAD sample. The statistics of the difference of the two group is calculated by Mann-Whitney test. **(D)** Box plot of the 32 CRC patients with the expression of genes that are expected with a direct protein-protein interaction (PPI). **(E)** Correlation plot for log expression measured by transcripts per million (TPM) for SLC7A11 and SLC3A2 based on COAD data (Log_2_(X)TPM; 461 cases). (**F)** Box plot from TCGA for COAD (461 cases) and Rectum adenocarcinoma (READ, 172 cases). Tumor and healthy are colored red and gray, respectively.

**Fig. 4B** shows a density plot for all 16 core-SLC7A11 genes. Some genes are very low expressed in CRC (SLC1A2), a few genes do not exhibit expression difference between tumor and normal tissue. Similar to SLC7A11 genes, the expression levels of most core genes are higher in tumor relative to normal samples. **Fig. 4C** confirms that the overall signature of all 16 listed genes remain highly significant for the difference of tumor relative to healthy tissue for COAD (Mann-Whitney p-value 1.9e- 67).

A small set of direct interacting proteins of SLC7A11 was compiled by UniProtKB and BioGrid. The confirmed direct interactors include SLC3A2 and CD44, but also KRTAP1-1, KRTAP1-3 and TTC30B. **Fig. 4D** analyzed the expression of major interacting genes from the CRC cohort (32 patients). Notably, the keratin-associated (KAP) family members and TTC30B levels of expression were too low, and these genes were not further analyzed. We report on the upregulated expression of direct interactors of SLC7A11 in tumor relative to healthy tissue **(Fig. 4D).** We further expanded the analysis to cover samples from TCGA (∼460 samples). A strong and significant correlation (R =0.41, p-value =5e-14) between SLC7A11 and SL3A2 expression at the individual level was confirmed (**Fig. 4E**).

These results suggest that the Xc- system (SLC7A11, SL3A2) is likely to be involved in the tumorigenesis process. Moreover, the upregulation of SLC7A11 in tumor samples was validated with high expression in patients with either COAD (461 cases) or READ (172 cases; **Fig. 4F**).

### Knowledge-based inspection of SLC7A11 determines its oncogenic potency

We sought to identify the network of SLCA11 correlated genes by considering gene perturbations in a cellular context. To this end, we tested the essentiality, specificity and efficacy of CRISPR dependency screens (see Methods). To further inspect the importance of SLC7A11 in colon cancer, we investigated the CRISPR-based dependency map for SLC7A11, SLC3A2 (and CD44) that comprises the Xc- system. Specifically, we compared genes that are most correlated following CRISPR-based gene depletion and focused on the overlapping genes displaying positively correlated signal (among the top 100 per each gene). **Fig. 5A** compared the genes that are most significantly recurrent in these CRISPR-Cas9 screening. The top co-dependent genes by CRISPR-Cas9 setting are expected to specify the degree of gene essentiality and replication fitness. There are 27 shared genes that are shared between the Xc- membrane transporters. We also observe a strong relatedness between the co-dependency genes of SLC7A5 and SLC3A2.

**Figure 5.**
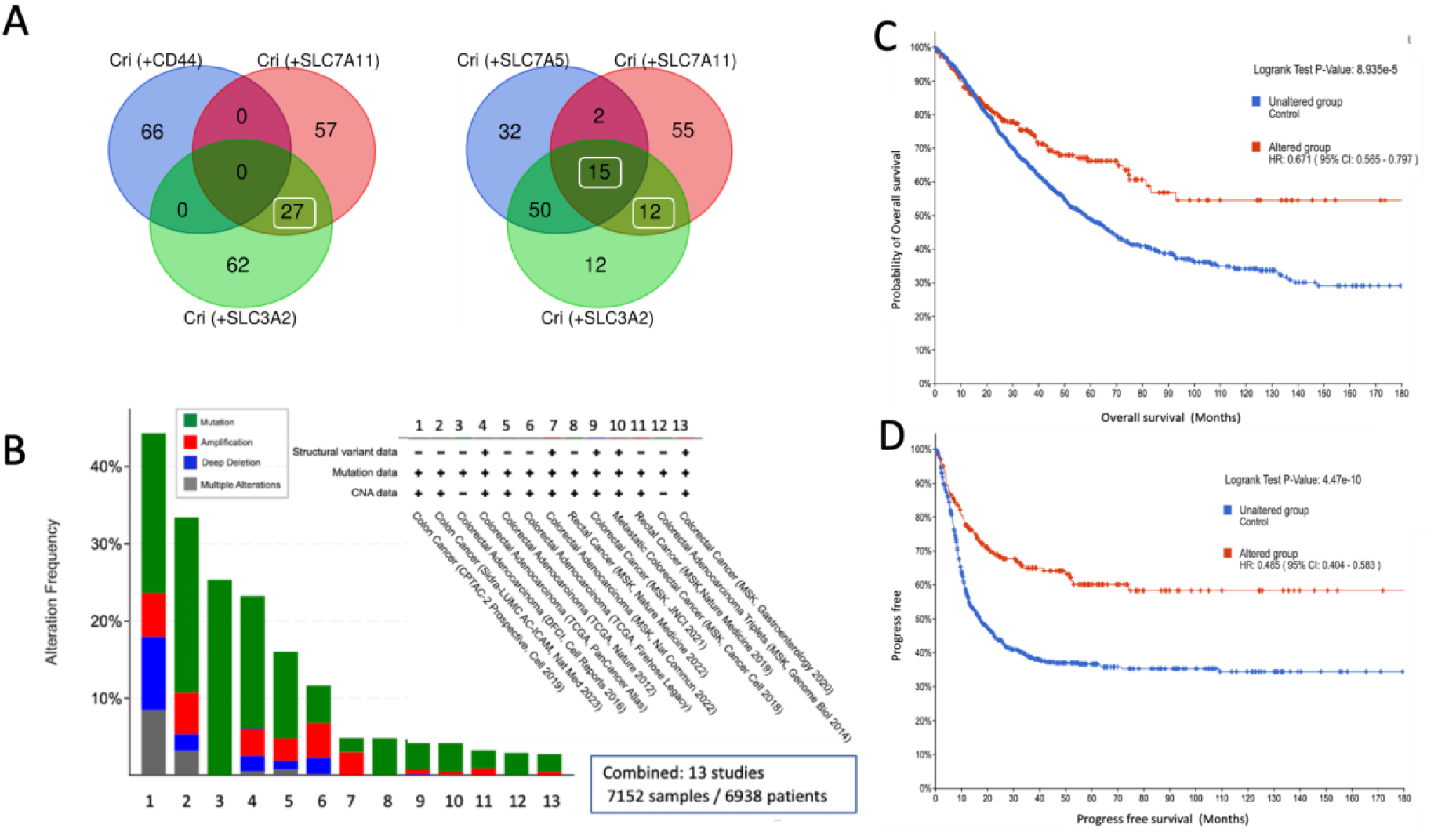
Gene set of CRISPR-induced Xc- fitness is consistent with patients’ CRC survival. **(A)** The analysis of top co-dependent genes by CRISPR-Cas9 setting (DepMap-based, see Methods). Only positively correlated genes from the 100 correlation-ranked genes list are included in the analysis. There are 27 shared genes (left, white frame) between the two membrane transporters and no genes overlapped with CD44. In contrast, the SLC7A5 displayed strong shared signal with SLC3A2 (Right, 65 genes) among them 15 genes are shared by all three gene sets. **(B)** Total 13 bowel cancer studies are listed according to the type of alteration in their genes (e.g., mutations, copy number variations; see legend for colors). We selected 13 of 19 studies reported in cBioPortal (a total of 7152 samples). All selected studies have at least 100 samples each, with the appendiceal cancer cohort excluded. **(C)** Probability of overall survival (OS) for 180 months is shown for affected set for the 27 shared genes (as in D). **(D)** Progress free survival (PFS) curve for 180 months. The survival plot indicates the unaltered and altered set (blue and red, respectively) for samples with alteration in any of the 27 shared genes. The statistical significance, Hazard ratio (HR) with 95% confidence is indicated.

For testing the impact of alteration in the overlapping genes (27) on the survival of CRC patients, we created a composed set from 13 independent studies (**Fig. 5B**) and analyzed the overall survival (OS, **Fig. 5C**) and progression-free survival (PFS, **Fig. 5D**). In both survival settings, the survival of the altered genes is enhanced. We observed that upon altering these key genes, a strong suppression in tumorigenesis is observed. The hazard ratio (HR) indicates an improved survival relative to the unaltered group (accounts for 90% of the samples). The HR for overall survival (OS) was 0.671 (**Fig. 5C**) and the progression free survival (PFS) was 0.485 (**Fig. 5D**). The genes when overexpressed in cancer cells support the progression of cancer and most likely resist process of apoptosis and other types of cell death.

### Cellular interpretation of overexpressed Xc- system in COAD

The observation that CRISPR co-dependent genes of the Xc- system resulted in strong clinical effect on survival, calls for identifying mechanistic explanation and biological pathways that connect Xc- system with cell growth and proliferation.

Among the 27 overlapping genes from the CRC cohort (32 patients, Supplementary **Table S2**), 6 genes blelong to the cation/anion and amino acid transporters of the SLC family (SLC29A1, SLC39A10, SLC5A6, SLC6A6, SLC7A11, and SLC7A5). The encoded proteins mediate transport across the cell membrane of specific metal ions, inorganic cations and anions, and amino acid. Additional upregulated gene products with ion-transporting potential belong to the ATPase family (e.g., ATP11A).

Among the overlapping co-dependent set (27 genes), the expression levels of 17 showed upregulation in the tumor samples. Inspecting this set shows that in addition to their role in amino acid transport (*e.g.,* the Xc- system), numerous representatives are enzymes that function in de novo biosynthesis of purine/pyrimidine nucleotides (FC >2; **Table 2**). We conclude that these genes specify the capacity of cancer cells to maintain high demand for protein synthesis (*e.g.,* amino acids), while exhibiting a strong signature for de novo nucleotide biosynthesis.

**Table 2.** Overlapping CRISPR co-dependent genes of Xc- system genes in COAD samples.

| Gene | Description | Main function | FC<br>T vs H<sup>a</sup> | FC<br>Met. vs<br>T | FC<br>Met. vs<br>H<sup>b</sup> |
| --- | --- | --- | --- | --- | --- |
| KRT16 | Keratin, type I cytoskeletal 16 | Intermediate filament | **13.13** | 0.13 | 1.73 |
| SLC7A11 | Cystine/glutamate transporter | Cystine transport as redox regulator | **9.22** | 0.34 | 3.15 |
| PRH2 | Proline rich protein HaeIII subfamily 2 | Secreted glycoprotein | **3.27** | 0 | 0 |
| MTHFD1 | Methylenetetrahydrofolate dehydrogenase | De novo Purine syntheses | **2.72** | 0.62 | 1.7 |
| ATIC | Formyltransferase/IMP cyclohydrolase | De novo purine biosynthetic pathway | **2.7** | 2.15 | **5.79** |
| RPIA | Ribose 5-phosphate isomerase A | Pentose-phosphate pathway | **2.65** | 0.77 | 2.03 |
| UMPS | Uridine monophosphate synthetase | De novo pyrimidine biosynthetic pathway | **2.65** | 6.34 | **16.79** |
| CAD | Transcarbamylase-dihydroorotase | de novo biosynthesis of pyrimidine nucleotides | **2.62** | 1 | 2.61 |
| TFRC | Transferrin receptor | cellular iron uptake | **2.51** | 1.91 | **4.79** |
| RAB36 | Ras related protein Rab-36 | Vesicle-mediated transport | **2.47** | 0.8 | 1.98 |
| SLC3A2 | 4F2 Cell surface antigen heavy chain | Transport of L-type amino acids | **2.27** | 0.54 | 1.22 |

<sup>a</sup>FC is the fold change. In bold face FC>2.0 for Tumor (T) vs healthy (H). <sup>b</sup>In bold face genes that amplified the metastatic (Met.) state.

**Table 2** shows that some of these genes do not only support tumorigenesis but actually contribute to the metastatic potential in COAD patients (marked by FC >1 for metastatic versus local tumor). The genes that showed such metastatic amplification are ATIC and UMPS, which act in nucleotide de novo synthesis, and TFRC, which allows iron uptake.

### Determinants of immune cell infiltration pattern in CRC

The tumor microenvironment (TME) is a crucial component in determining the response to immune checkpoint inhibitor (ICI) therapy. Resources were developed that allow assessment of tissue purity with respect to the presence of immune cell subtypes (compiled by TIME2.0; see Methods). The associations between cell fractions and treatment responses relied on considering progression-free survival (PFS) and overall survival (OS). Identifying distinct immune subgroups in CRC with varying responses to ICI therapy is a step toward achieving lasting ICI efficacy. For an unbiased approach, we tested the correlation of all 22 immune cell types with respect to the expression of SCLA711 and other genes that were highlighted in this study. For many of the cell types, statistical significance could not be achieved due to the low number of samples. All significant observations for COAD and READ are presented in Supplementary **Table S3**.

**Fig. 6** displays the scatter plot illustrating the relationship between infiltrates estimation and SLC7A11 expression. It shows the results for the tendency of the cancer samples to support immune cell infiltration in the case of COAD (**Fig. 6A**; 461 samples) and READ (**Fig. 6B**; 166 samples). The cofounding effect of purity is accounted for, and positive correlations are evident regarding CD8+ cells for COAD and READ samples. **Fig. 6C** shows a heatmap for two cell types: the hematopoietic stem cells and the myeloid-derived suppressor cells (MDSCs) across many cancer types. Importantly, MDSCs carry potent immunosuppressive activity and are closely associated with poor clinical outcomes in cancer (Tang et al. 2020). We conclude that among CRC patients (COAD and READ), there is a substantial positive correlation to CD8+ T-cells (**Figs. 6A-6B**) and MDSCs cells (**Fig. 6C**) that render the high expression of Xc- system. These observations raised a question regarding the success of immunotherapy in CDC patients.

**Figure 6.**
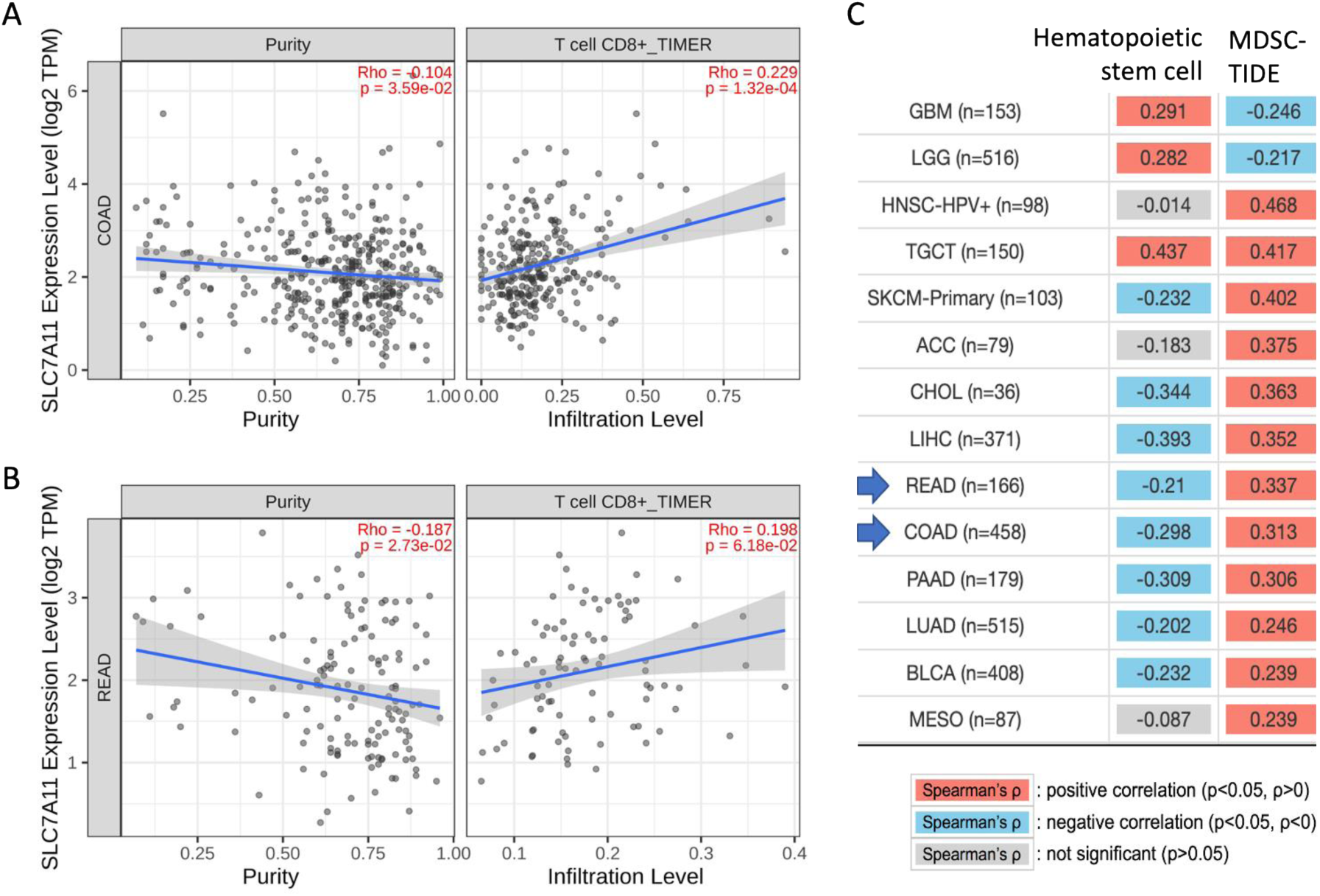
Correlation analysis of immune cell types and the data from TCGA for each individual. **(A)** The scatter plot showing the positive correlation between the abundance of CD8⁺ T cells and the expression of SLC7A11 in TCGA cohorts with 458 patients with COAD. The left panel indicates tumor purity, and the right panel indicates the infiltration of the T cells. **(B)** The scatter plot showing the positive correlation between the abundance of CD8⁺ T cells and the expression of SLC7A11 in TCGA cohorts with 166 patients for READ. The left panel indicates tumor purity, and the right panel indicates the infiltration of the T cells. The Spearman correlation test was based on TIMER2.0 algorithm. Tumor purity adjustment (Purity) was applied to account for the negative correlation of the SLC7A11 (Li et al. 2020). **(C)** Spearman correlation values (Rho) for a number of cell types are listed for hemopoietic stem cells and MDSCs. The p-values are adjustment by purity and positive and negative are marked by red and blue colors. Spearman p-values that are not significant are colored gray.

### Expression levels of Xc- system strongly correlate with COAD chemotherapy treatment

Due to the extreme degree of SLC7A11 upregulation (**Table 2**, FC of T vs H is 9.22) we challenged the prognostic capacity of the expression with respect to clinical treatments. RNA-seq data of 805 patients with COAD was therefore tested for their predictive capacity (marked by ROC p-value and AUC) for SLC7A11 and SLA3A2. **Fig. 7A** shows the predictive results for SLC7A11 with respect to all chemotherapy treatments that partitioned for responders (451 patients) and non-responders (354 patients). Although prediction power is rather limited (AUC =0.581) a higher mean expression level for non-responders was observed (p-value =3.7e-05). Opposite trend where a higher level of expression was observed for responders relative to non-responders was associated with SLC3A2 (AUC =0.577) **(Fig. 7B)**.

**Figure 7.**
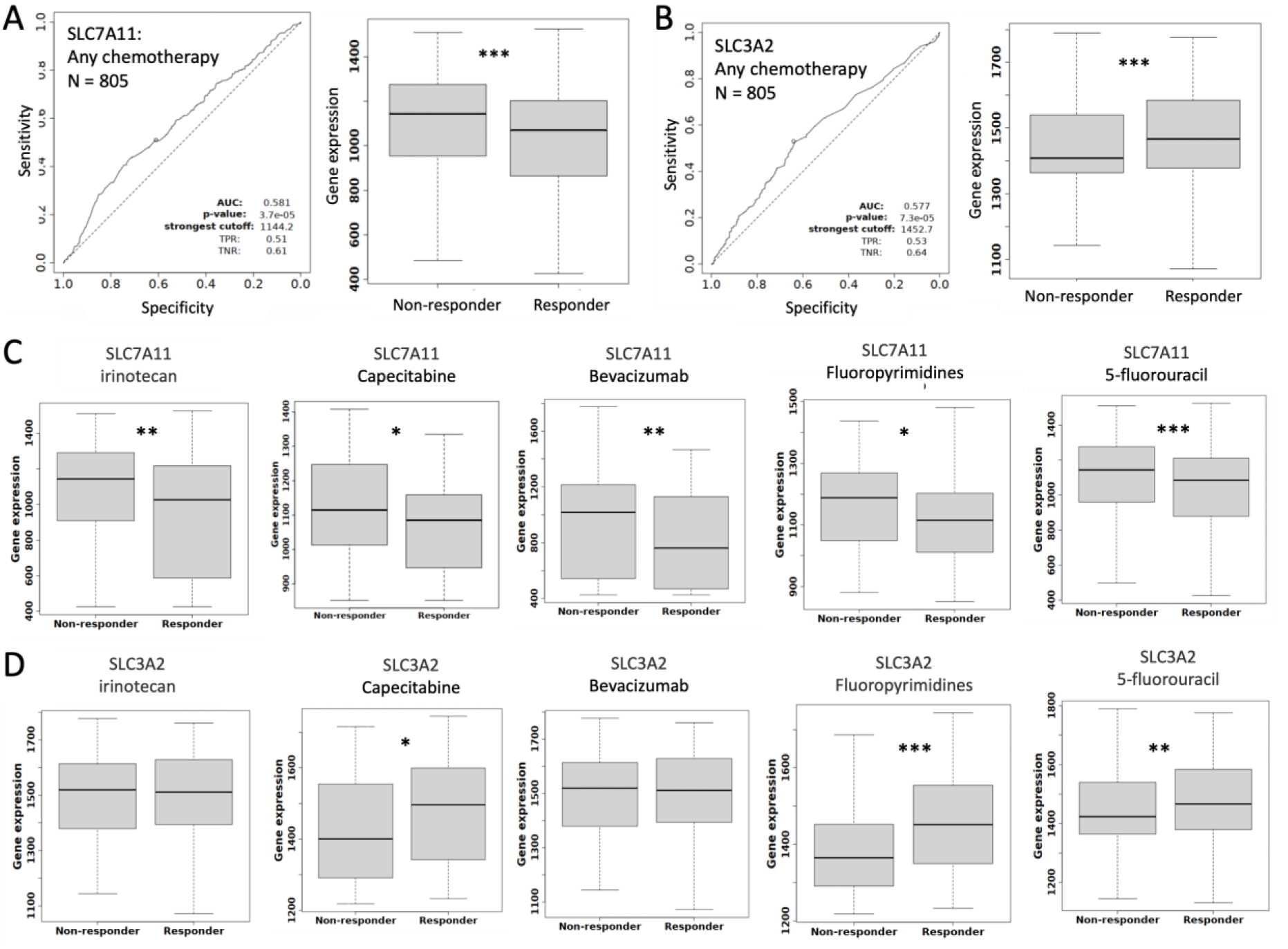
Expression levels of the genes of Xc- system by drug treatments and patients’ responsiveness. Analysis was performed by the KM-Plotted portal (KM-ROC). RNA-seq performed on 805 COAD patients partitioned by responders (451 patients) and non-responders (354 patients). **(A)** ROC-AUC and p-value are reported for SLC7A11 for all chemotherapies. **(B)** ROC-AUC and p-value are reported for SLA3A2 for all chemotherapies. **(C)** Box plot analysis of expression value with respect to responders and non-responders for specific reagents used in the CRC treatment for SLC7A11. **(D)** Box plot analysis of expression value with respect to responders and non-responders for specific reagents used in the CRC treatment for SLC3A2. The statistically significant marked by p-value at improving order of magnitude from <0.05 (*), <0.005 (**) to <0.0005 (***). Definition of a responder is by the RECIST criteria. We have not reported on results for treatment that applied to <100 patients.

The most common drug used in CRC patients is 5-flourouracil that includes 298 and 294 patients for the responders and non-responders, respectively. The partition in the box plots (**Fig. 7C**) shows that a slightly higher expression level for SLC7A11 was associated with the non-responders. Same trend (at a different confidence level) was observed for all other tested drugs in the relevant cohorts. The same analysis was applied to SLC3A2 which also showed a significant partition of responders and non-responders, in some but not all used drugs (**Fig. 7D**). Same analysis that was performed for oxaliplatin failed to reach any statistical significance (265 and 163 responders and non-responders, respectively).

Testing solid tumors identified a distinctive signature for SLC7A11 for the use of 5-fluorouracil with slightly higher expression for non-responder vs responder groups (294 and 298 patients, respectively) showing high significance (p-value of ROC =5.70e-04) but only a weak predictive power (AUC =0.576). Similar testing for alternative treatment such as oxaliplatin led to a border-line significance prediction potential (not shown). Only limited clinical data regarding the use of checkpoint inhibition therapy is available for CRC patients. Testing pretreatment by any immune checkpoint inhibitor therapy regarding SLC7A11 showed no signal for the success of treatment (including 533 responder and 570 non responders). We concluded that the information available is too limited to substantiate CRC patients’ stratification by their potential to successfully respond to T-cell-based immunotherapy.

## Discussion

In this study, we inspected the detailed molecular profiles of CRC patients through a cellular view combined with a knowledge-based network approach. For several genes, mechanistic relevance to cancer progression is presented to encourage further investigation. For example, among the strongest upregulated DEG, we observed several cytokines (i.e., CXCL1, CXCL2 and CXCL3; Fig. 3B). It may be a reflection of the abundance of MDSC cells which were positively correlated with the increased expression of SLC7A11 (Fig. 6C). It was shown that cancer-associated fibroblasts (CAF) via the secretion of chemokines (e.g., CXCL1, CXCL2) recruit myeloid cells to tumors and also drive the dysfunction of tumor-specific CD8^+^ T cells. A recent study proposed a prognostic model for CRC that is based on the immune cell composition in the tumor samples (Ye et al. 2019). With the effort to stratify CRC patients for improved clinical management and outcomes, immune cell expression signatures were classified into four subtypes that aim to capture the degree of T-cell dysfunction and exclusion (Tang et al. 2020; Zhang et al. 2020). While the immune classification of cancers is of utmost importance for prognostics, as predictive factors for chemotherapies and immune checkpoint inhibitor therapy, current knowledge remains inconsistent across different datasets (e.g., across 15 CDC datasets compiled in TIDE resource (Fu et al. 2020)).

Among the unregulated genes in COAD, Transferrin receptor (TFRC) was found to be even more enhanced in metastatic tumors (Table 2), suggesting an important role for iron uptake in these cells. In agreement with our observation, TFRC was shown to be an essential factor in nucleotide biosynthesis, DNA repair, and cell survival based on its crucial role in iron accumulation and ensuing iron-dependent activation to maintain the nucleotide pool and sustain proliferation in colorectal tumors (Schwartz et al. 2021). Congruently, TFRC was recently suggested as an attractive target for inhibiting tumor growth, as reduction of iron influx can lead to DNA damage and apoptosis (Kim et al. 2023).

We have focused on the role of the Xc- system in CRC cancer and showed that it acts as a hub connecting the elaborate strong signature of mitotic cells and cell cycle with the metabolic program (Fig. 2B). SLC7A11 may have opposite effects in different cancer cells. For example, in a glucose-deprivation state (as in glioblastoma) the overexpression of the Xc- system induces oxidative stress and apoptosis. In contrast, it is a strong mediator for cell viability in CRC and other cancer types. In such cases, the suppression of SLC7A11 function (*e.g.,* by p53 or BECN1) can activate ferroptosis which makes the tumor sensitive to radiotherapy. To further analyze the cellular role of Xc-, we inspected the downstream glutathione pathways (e.g., GPX4, GPX8). We observed that the expression of GPX genes were unchanged within the cohort of 32 patients (Supplementary Table S1), with no co-dependency in the CRISPR screening results. Essential and co-essential genes across many cell lines are a useful approach to identify shared pathways (Arnold et al. 2022). Using the DepMap platform, we analyzed GPX4 and GPX8 expression levels with respect to the effect size of CRISPR-based SLC7A11 knockdown across 39 cells originated from COAD. The GPX4 and GPX8 genes had negative Spearmen correlation (R: -0.449 and -0.205, respectively), suggesting the involvement of alternative pathways in the Xc- tumorigenesis.

Cellular models successfully used to identify overlooked pathways for SLC7A11 dependent ferroptosis. For example, using colon cancer cell lines (HCT116, LoVo, and HT29) confirmed a regulatory loop between PERK and SLC7A11 through transcription factor ATF4 (Saini et al. 2023). The suppression of PERK or ATF4 reduces SLC7A11 expression in these cells. A side benefit of our study was to highlight experimental cell lines that were most appropriate for future functional assays. Specifically, for the minimal set of shared genes of the SLC transporters (Fig. 5A) we highlighted 15 overlapping genes. A number of these genes (CAD, MTHFD1, UMPS, SDHB and SLC7A5 itself) were also listed among the most affected genes by CRISPR screen, and especially in COLO-205 metastatic COAD cell line. These essential genes exhibited a strong effect on loss of fitness (see Methods) and therefore are attractive targets for drug testings.

We demonstrated that the expression level of SLC7A11 can (slightly) predict chemotherapy responsiveness (Fig. 7). It is thus beneficial to apply anticancer therapies that downregulated the levels of the Xc- system directly (e.g., by SLC7A11 blockers) or indirectly. Several small molecules and inhibitors targeting SLC7A11 have been developed and are being investigated for their therapeutic potential (Xu et al. 2020). An indirect pathway impacting Xc- system involves the use of PD-L1 blockade therapy that leads to an increase lipid ROS production, and through STAT1 attenuates SLC7A11 expression. More mediators include JAK and ATM (following radiotherapy) and other studies focus on the strong link of SLC7A11 and ferroptosis in CRC. It was shown that the loss of PERK (Saini et al. 2023) or vitamin D (Guo et al. 2023) promoted downregulation of SLC7A11 and consequently induced ferroptosis.

Our study further emphasizes the importance of developing targeted therapies that rely on understanding the link of SLC7A11 to ferroptosis while utilizing the nutrient dependency of cancer cells. Overall, targeting SLC7A11 by multiple routes can be an effective strategy to enhance therapeutic efficacy, induce regulated cell death and ultimately improve patients’ outcomes. While we focused mostly on the levels of gene expression and cellular response, other regulatory mechanisms that affect SLC7A11, including the cellular network of post-transcriptional controls by microRNA (miRNA), or the unexplored landscape of post translational modifications (PTMs) in tumor samples, remain interesting strategies that need to be further elucidated.

## Conflicts of Interests

The authors declare that they have no conflicts of interest to report regarding the present study.

## Ethics approval and consent to participate

The Ethics Committee of University Magdeburg approved this study

## Abbreviations

AUC: area under the curve
CAF: cancer-associated fibroblasts
CD44v: CD44 variant
CHIP-seq: chromatin Immunoprecipitation Sequencing
COAD: colorectal adenocarcinoma
CPM: counts per million
CRC: colorectal cancer
DEG: differential expression genes
FC: fold change GO gene ontology
GPX: glutathione peroxidases
GSH: glutathione
GTEx: genotype tissue expression
ICI: immune checkpoint inhibitor
OS: overall survival
PCA: principal component analysis
PFS: progression-free survival
READ: rectum adenocarcinoma
RNA-seq: RNA sequencing
ROS: reactive oxygen species
TCGA: the cancer genome atlas
TME: tumor microenvironment
TMM: trimmed mean of the M-values
TPM: transcript per million
xCT: solute carrier family 7, member 11

